# The preference of a fragrance created by a perfumer is associated with one single-nucleotide polymorphism in OR5A1 gene

**DOI:** 10.64898/2026.01.14.699479

**Authors:** Marylou Mantel, Morgane Dantec, Pierre Arnold, Lenka Tisseyre, Veronica Pereda-Loth, Manon Gabriel, Denis Pierron, Moustafa Bensafi

**Author notes:** co-first authors / (Marylou Mantel, Morgane Dantec, Pierre Arnold). co-last authors (Denis Pierron, Moustafa Bensafi).

## Abstract

From ancient incense to Chanel No. 5, perfumes are almost ubiquitous across the world and across history yet highly diverse. While they reflect the artistry of the creator and the personality of the wearer, the neurobiological processes driving fragrance preference are unknown.

Recent research demonstrated that genetic variability in olfactory receptors modulates odor perception; however, regarding the preference for complex real-world stimuli, the relative importance of peripheral sensory encoding versus top-down cognitive evaluation—shaped by individual life history—remains a subject of debate.

Here, we evaluate whether the perception of a commercially available fragrance depends on genetics, using the example of beta-ionone, a molecule routinely used in perfume composition, whose perception and appreciation have been linked to a single-nucleotide polymorphism in the *OR5A1* gene (*rs691536*). A total of 168 participants rated their preference for three versions of a perfume designed by a professional perfumer, differing only in the concentration of β-ionone used in the formula.

Results showed that carriers of the G allele rated the 0% β-ionone fragrance significantly higher than the 10% and 50% versions, while AA homozygotes showed no discrimination between the perfumes. By focusing on just one SNP (among hundreds of olfactory genes) and a single molecule, this study offers a proof of concept regarding the influence of genetics on aesthetic fragrance appreciation, advocating for the integration of genomic data to better understand the global diversity of perfume preferences.

## RESULTS

Better access to genome-wide association studies has led to growing investigation of the genetic basis of single-molecule olfactory perception^1^. However, it is not yet clear if and how genetic background might change the way we perceive and appreciate everyday olfactory objects such as perfumes, even though a number of these single molecules are routinely used in the industry. One of them is β-ionone, a naturally occurring compound with a violet-like odor regularly used in perfume formulation^2,3^. The detection threshold of this compound, as well as its pleasantness, is strongly associated with a polymorphism in the OR5A1 gene, which may also impact the pleasantness of foods artificially spiked with β-ionone^4,5^.

Here, we aimed to extend the question of the influence of variability in the OR5A1 gene to the appreciation of more ecological stimuli such as perfumes. To this end, we analyzed the impact of β-ionone perception variability on the hedonic perception of a commercially distributed perfume by collaborating with its French creator to develop three distinct variations containing concentrations of 0%, 10%, and 50%.” In total, 168 participants included across three different French cities (Toulouse, Lyon and Reims) were asked to rate and rank these three versions according to preference, and their answers were linked to genotype for OR5A1. This allowed for the testing of two hypotheses: 1/ people carrying at least one copy of the functional G allele of OR5A1 will have different hedonic preferences among perfumes containing various amounts of β-ionone compared to people with only non-functional allele A, and 2/ more specifically, we can expect that people carrying at least one copy of the functional G allele of OR5A1 will be preferring more the perfumes containing less β-ionone compared to people with only non-functional allele A, in line with previous results^4^.

### Description of the genotype groups

Among the 168 participants, 71 (42%) carried the AA genotype at the *rs691536* location for the OR5A1 gene, 81 (48%) the AG genotype and 16 (10%) the GG genotype. As Jaeger et al’s^4^ results showed that the phenotypic difference is related to the presence of at least one dominant G allele, AG and GG genotype groups were subsequently combined for the analysis. Regarding the demographic characteristics of each genetic group, there was no significant difference in age (W = 3247, p = .53) nor in sex ratio (X^2^(1) = 0.09, p = .76).

### Validation of the correspondence between genetic variation in OR5A1 and sensitivity to β-ionone

In order to first verify the association between genotype and β-ionone thresholds, a subset of 98 participants were administered a β-ionone threshold test, there was a significant difference in thresholds according to genotype (W= 2315, p < .001): indeed, people with genotype AA (n= 46) had a mean threshold of 4.6_∓_2.1, equating to around 280 ppb, compared to 13.2_∓_3.4 for people with genotype AG-GG (n= 52), equating to around 0.7 ppb (see *Supplementary Figure S1*). Participants with genotype AG-GG thus had a 400 times higher sensitivity to β-ionone compared to the ones with genotype AA.

Regarding demographic characteristics, we did not find a significant association between β-ionone thresholds and age (r_S_(98) = -.10, p = .29), nor sex of the participants (W = 1062, p = .93).

### Association between genetic variation in OR5A1 and preference for perfumes containing β-ionone

The main analysis showed a significant association between genotype and perfume preference (X^2^(2) = 8.19, p = .017). More specifically, people with the AG/GG genotype group preferred more the sample with 0% β-ionone compared to the other samples, whereas there was no difference in preference for individuals from the AA group (p_Holm_= .026), as shown in *Figure 1*.

**Figure 1.**
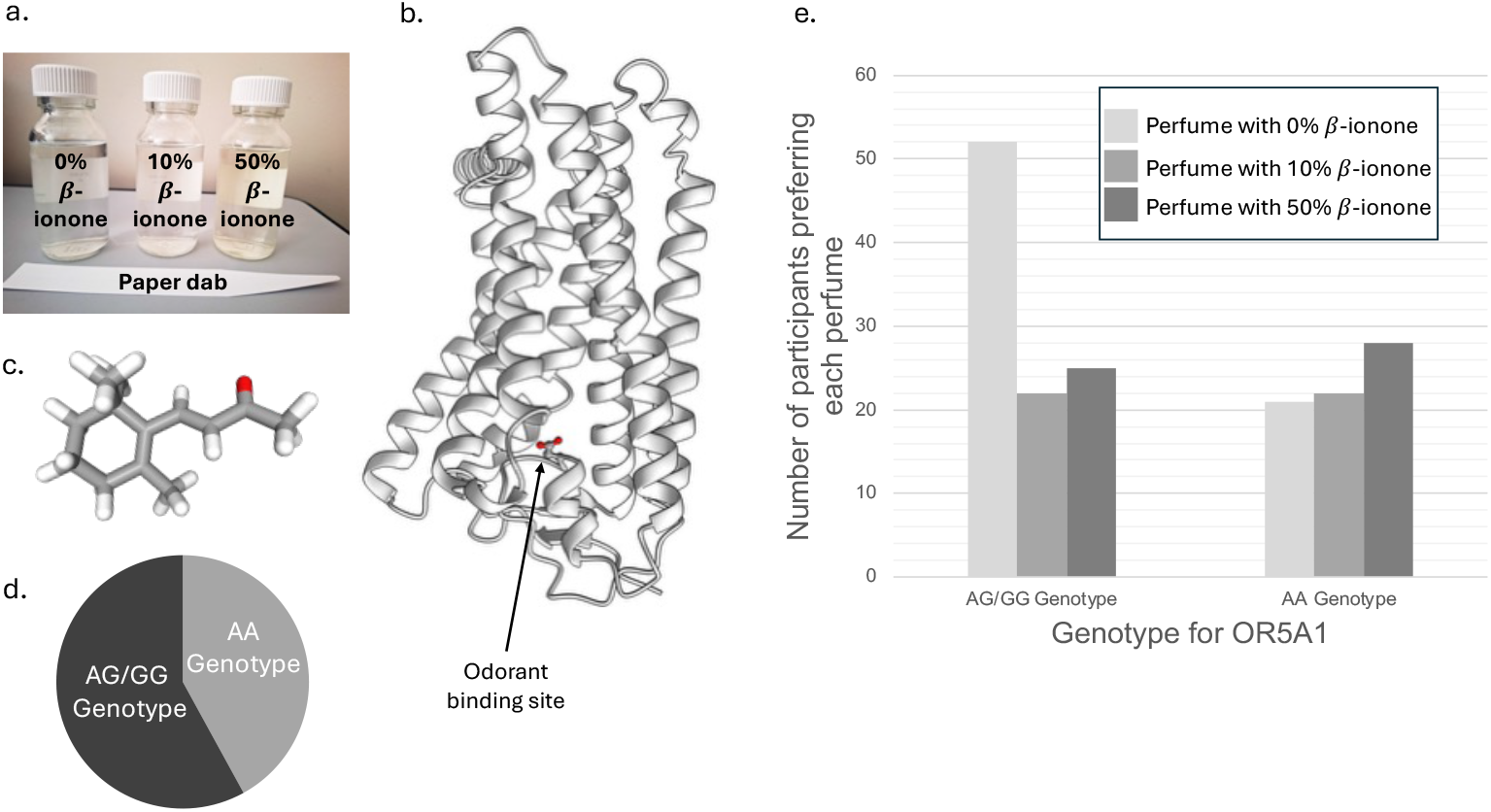
Summary of the methodology and main results of the experiment. a. Picture of the three perfume stimuli and paper dabs used in the study. b. Proportion of individuals in each genetic group, either AA or AG/GG genotype for the rs691536 location on the OR5A1 gene. c. 3D structure of OR541, with the odorant binding site represented in red whose structure is affected by the rs691536 single-nucleotide polymorphism. d. Graphical representation of the number of participants preferring each of the three perfumes depending on genotype group.

Regarding individual perfume ratings (see *Supplementary Figure S2*), there was a significant interaction between genotype and β-ionone concentration on pleasantness evaluations (F(2,102) = 4.16, p = .02). Indeed, the perfume containing 0% β-ionone was rated as more pleasant than the 50% one for people with the AG/GG genotype, but not for the participants with the AA genotype (p_Holm_ = 0.01). For intensity, there also was a significant interaction effect (F(2,99) = 3.66, p = .029), with people with genotype AG/GG rating the perfume containing 0% β-ionone as stronger than the 50% one, but not people with genotype AA (p_Holm_ = 0.04).

### Other individual factors affecting preference for perfumes containing β-ionone

In order to explore other common demographic factors that may influence preference in individuals, a multinomial logistic regression was conducted on perfume preference with genotype group, sex and age as predictors. The model was statistically significant (X^2^(6) = 17.3, p = .008, R^2^ = .05) and age and genotype, but not sex, were significant predictors in the different preferences for 0%, 10% or 50% β-ionone perfumes. More specifically, there was a higher likelihood in preferring the sample with 0% β-ionone over the one with 10% for people that were carrying genotype AG or GG (B = 0.95, p = .020, CI [0.15, 1.75]) and who were younger (B = 0.043, p = .027, 95% CI [0.0048, 0.08]). There was also a higher likelihood in preferring the sample with 0% β-ionone over the 50% one for people that were carrying genotype AG or GG ((B = 1.04, p = .007, 95% CI [0.29, 1.79]) and younger ((B = 0.031, p = .098, 95% CI [-0.0058, 0.070]), although this last association was tendential only.

However, these results must be mitigated by two elements. First, the sample was quite young (median age = 24, IQR = 10) and age was not normally distributed. Second, data was collected in three different sites, with age differences between the samples (X^2^(2) = 25.2, p < .001).

## Discussion

In light of the importance of perfumes, both across the world and across history, as well as their relationship with personal taste, creativity and individuality, the question of what influences perfume appreciation and choice is relevant. Consumer research emphasizes the importance of factors like marketing or packaging^6^, but also certain personality traits and social conformity ^7^. The appreciation of the fragrance itself may be related to factors already known to influence odor perception and pleasantness in general: age, gender, previous experiences and cultural background, but also more physiological dimensions such as hunger state, menstrual cycle and certain diseases^8^. However, the contribution of genetics, through variations in olfactory receptors structure and functionality, is less understood. Although the perception and hedonic value of single molecules have been associated to single-nucleotide polymorphisms^1,9,10^, whether evaluation of complex odorant mixtures like perfumes is also impacted has not yet been demonstrated.

In line with our hypotheses, the results of this study showed different preferences for perfumes containing β-ionone according to genotype: participants with genotype AG/AA preferred more often the perfume without the molecule compared to the ones with either 10% or 50%, whereas there was no difference in preference for participants with genotype AA. Regarding the individual ratings of the perfumes, participants with genotype AG/AA found that the sample without β-ionone was both more pleasant and less intense compared to the sample with 50%, while no significant difference was found for the other genotype group.

Such a preference for stimuli without β-ionone for highly sensitive people is congruent with previous work^4,5^. Jaeger et al^4^ reported a difference in hedonic appreciation of β-ionone in isolation: people with genotype AG/GG and highly sensitive to the molecule tend to like it less compared to less sensitive people with genotype AA and use different qualitative descriptors to qualify it. The same team also investigated the differences in liking foods spiked with β-ionone, such as chocolate or apple juice, and showed that highly sensitive participants with genotype AG/GG liked less the samples containing the molecule compared to the unspiked ones, whereas less sensitive participants with genotype AA did not differ in their ratings. However, it is interesting to note that, contrary to previously reported intensity ratings of β-ionone in isolation in which the stronger the concentration of β-ionone, the stronger the intensity of the odor, the present study showed that highly sensitive individuals perceived the perfume without the molecule as the most intense one. This could be due to a variety of factors, including perceptual interactions with the others components of the mixture^11^ or inhibition processes between olfactory receptors^12^.

Overall, the impact of β-ionone sensitivity on the processing of odor mixtures is likely very complex, due to interactions with other compounds and other polymorphisms in olfactory receptors. Indeed, recent work suggests that the molecule acts as an antagonist for the processing of other molecules like methanethiol^13^. Also, genetic differences in sensitivity to other molecules commonly used in fragrances have been reported. For example, detection of cis-3-hexenol, a compound with a fresh, grassy odor found in several foods and used in cosmetic formulation, is partly influenced by a polymorphism in the OR11H7P gene^10^. The perception of musk, a family routinely used in the fragrance industry, is affected by a polymorphism in the OR5AN1 gene as well^14^. Regarding the latter, a polymorphism in the OR5AN1 gene linked to lower macrocyclic musk detection is associated with the AG/GG genotype for OR5A1, suggesting that people who are more sensitive to β-ionone are also less sensitive to musk^14^.

In conclusion, this study is the first to bring to light the direct association between a single-nucleotide polymorphism for a particular olfactory receptor and the appreciation of a real perfume. Although further work is necessary to better understand the complex interaction between genetic variability and other known factors in the perception and pleasantness of complex odorant mixtures, the present results emphasize the contribution of genetics in fundamental and consumer research on perfume preference.

## Supporting information

Supplementary Results

## RESOURCE AVAILABILITY

### Lead contact

Requests for further information and resources should be directed to and will be fulfilled by the lead contact, Marylou Mantel.

### Materials availability

This study did not generate new unique reagents.

### Data and code availability

All data reported in this paper will be shared by the lead contact upon request. This paper does not report original code. Any additional information required to reanalyze the data reported in this paper is available from the lead contact upon request.

## ACKNOWLEDGMENTS

This work was supported by a grant from the Per Fumum foundation, a non-profit organization promoting research on olfaction and perfume.

## AUTHOR CONTRIBUTIONS

Conceptualization: M.B., D.P., M.M., V. P-L.; Methodology: M.B., D.P., M.M., M.D. P.A., V. P-L.; Formal analysis: L.T., M.M., M.D. P.A.; Investigation: M.M., M.D. P.A., M.G.; Writing-original draft; M.M.; Writing-review and editing: M.M., M.D., P.A., M.B., D.P.; Supervision: M.B., D.P.; Funding acquisition: M.B., D.P., M.M

## STAR Methods

## STUDY PARTICIPANTS DETAILS

A total of 171 participants (71% women, 15-60 years old, mean age 28.2_∓_10.6) was recruited on university and research campuses in Toulouse, France between April and July 2023, in Lyon, France, in March 2024, and in Reims, France, between February and July 2025. Inclusion criteria were the following: age 15 and older, no known acute or chronic olfactory disorder, no known acute or chronic gustatory disorder, no known food allergy. All participants gave their written and oral consent for the tests and DNA sampling. The study was conducted according to the Declaration of Helsinki and approved by a French ethical committee (*CPP Sud-Méditerranée II - 2022-A00865-38*). Participants were not financially compensated but received a snack at the end of the experiment.

## METHOD DETAILS

The three olfactory stimuli used in the study consisted of three variations of the same perfume created by a perfumer. The three mixtures comprised 50% of dipropylene glycol (DPG, solvent) and β-ionone pair (*DPG/β-ionone ratio mixture 1: 50%/0%; mixture 2: 40%/10%; mixture 3: 0%/50%*), and 50% other compounds including hedione, linalool, phenylethanol, musk, helvetolide, geranyl acetate, citrol, and cedramber as the main constituents. The three perfume samples were compounded and conditioned in 50ml clear glass lab bottles at the same time and stored at ambient temperature. At the time of the experiment, the perfumes were presented to the participants on standard paper labs dipped in the bottles.

After an introduction and explanation of the study, participants were presented with three perfume labs dipped in the three perfume stimuli (either containing 0%, 10% or 50% β-ionone in their formula) in a counterbalanced order. In a first step, they were asked to rate the degree of similarity between each perfume by comparing them in pairs, using a Likert scale from 1 (not similar at all) to 9 (identical). Then, they had to rank the three perfumes from the most liked to the least liked. In a second step, individuals were asked to rate each perfume on several parameters, again using a Likert scale from 1 (not at all) to 9 (extremely): degree of pleasantness, of intensity, of irritation and of familiarity. They also had a space to freely describe the perfume in their own words.

A threshold detection test to β-ionone (4-(2,6,6-trimethyl-1-cyclohexenyl)-3-buten-2-one, CAS = 14901-07-6) was administered to 100 participants, adapted from the TDI test. Concentrations were chosen according to previous studies investigating perceptual detection threshold for β-ionone^15,16^. The test included 16 dilution steps, starting from 3000 ppb and halved at each step, up until step 16 corresponding to 0.1 ppb. Each dilution step was conditioned with 4mL of distilled water, in opaque 15mL glass vials, and was coupled with 2 unscented vials also containing only water. The test was administered following the staircase method of the Sniffing Sticks ^17^.

Another psychometric test was also administered to the participants to control their global olfactory capabilities, in the form of a shortened version of the European Test for Olfactory Capabilites ^18,19^. After this second test, participants gave a saliva sample for genotyping. The single nucleotide polymorphism (SNP) of interest *rs691536* was genotyped for all individuals. Participants were then categorized into two groups based on their allele combination for this SNP: individuals with an AG or GG genotype versus individuals with an AA genotype. As genotyping was inconclusive for 3 out the 171 participants, the subsequent statistical analyses were conducted on the remaining 168 individuals.

At the end of the experimental session, participants also filled in a questionnaire, including demographic items and elements regarding their food habits.

## QUANTIFICATION AND STATISTICAL ANALYSIS

Data was organized and analyzed with the RStudio software (version 2025.09.0+387) as well as the JAMOVI and JASP software. Non-parametric tests were used as data were not normally distributed. For the analysis of variance with two factors, two-way mixed ANOVAS for trimmed means were conducted, as suggested by Wilcox (2012), using the WRS2 package (version 1.1-6).

## References

1. Trimmer, C., Keller, A., Murphy, N.R., Snyder, L.L., Willer, J.R., Nagai, M.H., Katsanis, N., Vosshall, L.B., Matsunami, H., and Mainland, J.D. (2019). Genetic variation across the human olfactory receptor repertoire alters odor perception. Proc. Natl. Acad. Sci. U. S. A. 116, 9475–9480. 10.1073/pnas.1804106115.

2. Lalko, J., Lapczynski, A., McGinty, D., Bhatia, S., Letizia, C.S., and Api, A.M. (2007). Fragrance material review on β-ionone. Food Chem. Toxicol. 45, S241–S247. 10.1016/j.fct.2007.09.052.

3. Paparella, A., Shaltiel-Harpaza, L., Ibdah, M., Paparella, A., Shaltiel-Harpaza, L., and Ibdah, M. (2021). β-Ionone: Its Occurrence and Biological Function and Metabolic Engineering. Plants 10. 10.3390/plants10040754.

4. Jaeger, S.R., McRae, J.F., Bava, C.M., Beresford, M.K., Hunter, D., Jia, Y., Chheang, S.L., Jin, D., Peng, M., Gamble, J.C., et al. (2013). A Mendelian Trait for Olfactory Sensitivity Affects Odor Experience and Food Selection. Curr. Biol. 23, 1601–1605. 10.1016/j.cub.2013.07.030.

5. Jaeger, S.R., Reinbach, H.C., Roigard, C.M., McRae, J.F., Pineau, B., Chheang, S.L., Beresford, M.K., Rouse, S.A., Jin, D., Paisley, A.G., et al. (2014). Sensory characterisation of food and beverage stimuli containing β-ionone and differences between individuals by genotype for rs6591536. Food Res. Int. 62, 205–214. 10.1016/j.foodres.2014.02.038.

6. Saed, R.A., Salih, M.A., Hussien, A.H., and Swwedan, N. (2022). The impact of perfume packaging on consumer buying behaviour of Jordanian females. Int. J. Bus. Excell.

7. Ou, C.-C., and Chuang, H.-H. (2023). Exploring the Factors that Influence Consumers to Purchase Perfume Products. Int. J. Prof. Bus. Rev. Int J Prof Bus Rev 8, 18.

8. Rouby, C., Pouliot, S., and Bensafi, M. (2009). Odor hedonics and their modulators. Food Qual. Prefer. 20, 545–549.

9. Keller, A., Zhuang, H., Chi, Q., Vosshall, L.B., and Matsunami, H. (2007). Genetic variation in a human odorant receptor alters odour perception. Nature 449, 468–472.

10. McRae, J.F., Mainland, J.D., Jaeger, S.R., Adipietro, K.A., Matsunami, H., and Newcomb, R.D. (2012). Genetic variation in the odorant receptor OR2J3 is associated with the ability to detect the “grassy” smelling odor, cis-3-hexen-1-ol. Chem. Senses 37, 585–593. 10.1093/chemse/bjs049.

11. Thomas-Danguin, T., Sinding, C., Romagny, S., El Mountassir, F., Atanasova, B., Le Berre, E., Le Bon, A.-M., and Coureaud, G. (2014). The perception of odor objects in everyday life: a review on the processing of odor mixtures. Front. Psychol. 5. 10.3389/fpsyg.2014.00504.

12. Pfister, P., Smith, B.C., Evans, B.J., Brann, J.H., Trimmer, C., Sheikh, M., Arroyave, R., Reddy, G., Jeong, H.-Y., Raps, D.A., et al. (2020). Odorant Receptor Inhibition Is Fundamental to Odor Encoding. Curr. Biol. 30, 2574–2587.e6. 10.1016/j.cub.2020.04.086.

13. Fukutani, Y., Abe, M., Saito, H., Eguchi, R., Tazawa, T., de March, C.A., Yohda, M., and Matsunami, H. (2023). Antagonistic interactions between odorants alter human odor perception. Curr. Biol. CB 33, 2235–2245.e4. 10.1016/j.cub.2023.04.072.

14. Sato-Akuhara, N., Trimmer, C., Keller, A., Niimura, Y., Shirasu, M., Mainland, J.D., and Touhara, K. (2023). Genetic variation in the human olfactory receptor OR5AN1 associates with the perception of musks. Chem. Senses 48, bjac037. 10.1093/chemse/bjac037.

15. Plotto, A., Barnes, K. w., and Goodner, K. l. (2006). Specific Anosmia Observed for β-Ionone, but not for α-Ionone: Significance for Flavor Research. J. Food Sci. 71, S401–S406. 10.1111/j.1750-3841.2006.00047.x.

16. Jaeger, S.R., McRae, J.F., Bava, C.M., Beresford, M.K., Hunter, D., Jia, Y., Chheang, S.L., Jin, D., Peng, M., Gamble, J.C., et al. (2013). A Mendelian Trait for Olfactory Sensitivity Affects Odor Experience and Food Selection. Curr. Biol. 23, 1601–1605. 10.1016/j.cub.2013.07.030.

17. Hummel, T., Sekinger, B., Wolf, S.R., Pauli, E., and Kobal, G. (1997). “Sniffin” sticks’: olfactory performance assessed by the combined testing of odor identification, odor discrimination and olfactory threshold. Chem. Senses 22, 39–52. 10.1093/chemse/22.1.39.

18. Thomas-Danguin, T., Rouby, C., Sicard, G., Vigouroux, M., Farget, V., Johanson, A., Bengtzon, A., Hall, G., Ormel, W., De Graaf, C., et al. (2003). Development of the ETOC: a European test of olfactory capabilities. Rhinology 41, 142–151.

19. Joussain, P., Bessy, M., Faure, F., Bellil, D., Landis, B.N., Hugentobler, M., Tuorila, H., Mustonen, S., Vento, S.I., Delphin-Combe, F., et al. (2016). Application of the European Test of Olfactory Capabilities in patients with olfactory impairment. Eur. Arch. Otorhinolaryngol. 273, 381–390. 10.1007/s00405-015-3536-6.

20. Wilcox, R.R. (2012). Introduction to robust estimation and hypothesis testing 3rd ed. (Academic Press).

