## Supplementary Results for "The preference of a fragrance created by a perfumer is associated with one single-nucleotide polymorphism in OR5A1 gene"

**Supplementary materials**

*Figure S1. Distribution of measured thresholds for beta-ionone depending on genotype group.*

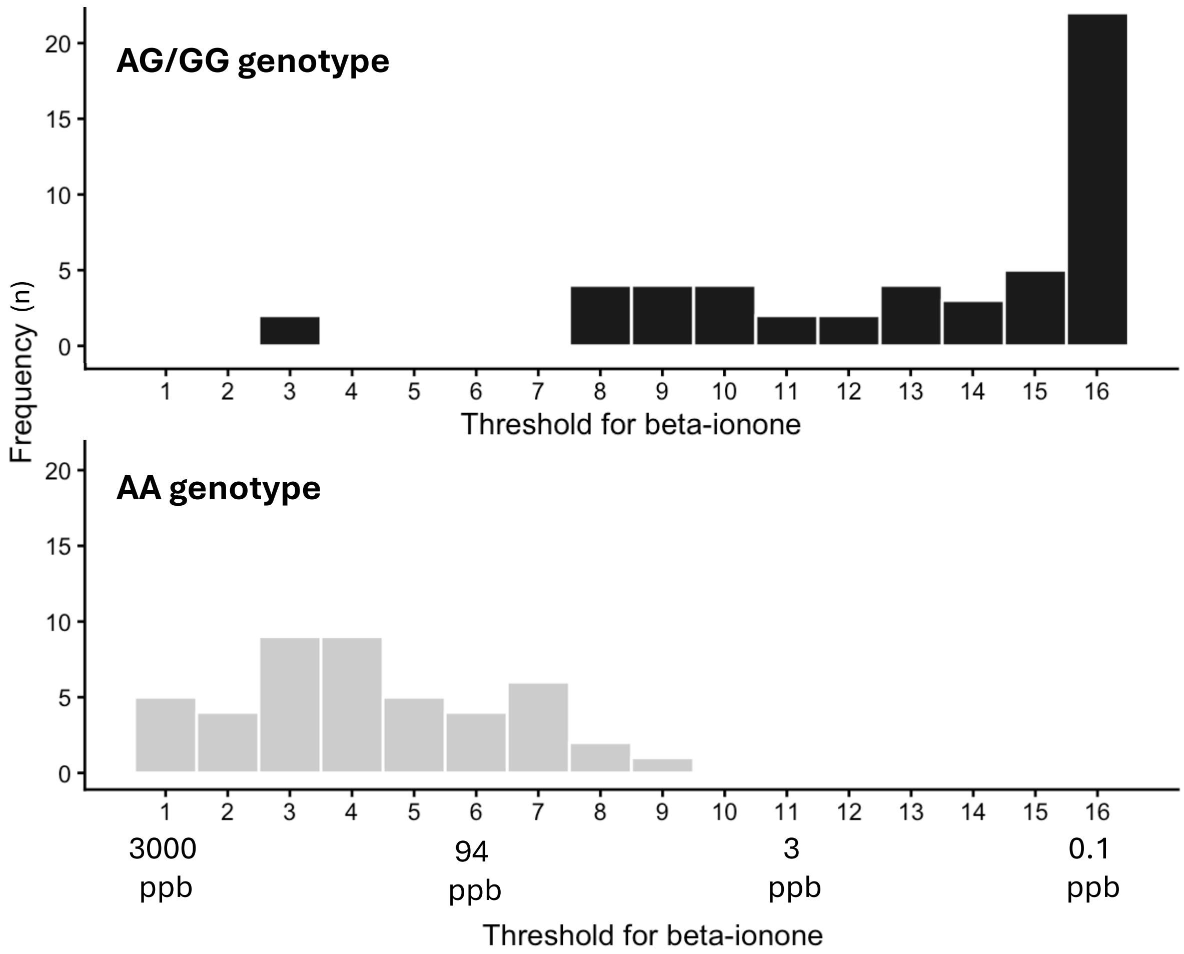


*Figure S2. Mean perceptual ratings for the three perfumes depending on genotype group. a. Hedonic ratings. b. Familiarity ratings. c. Intensity ratings. d. Irritation ratings. * p < .05, ** < .01. Error bars correspond to standard error.*

The perfume containing 0% β-ionone was rated as more pleasant than the 50% one for people with the AG/GG genotype, but not for the participants with the AA genotype ($p_{Holm}$ = 0.01). For intensity, there also was a significant interaction effect ($F(2,99) = 3.66$, $p = .029$), with people with genotype AG/GG rating the perfume containing 0% β-ionone as stronger than the 50% one, but not people with genotype AA ($p_{Holm}$ = 0.04). There was no significant effect of concentration, genotype nor interaction effect for the familiarity and irritation ratings.

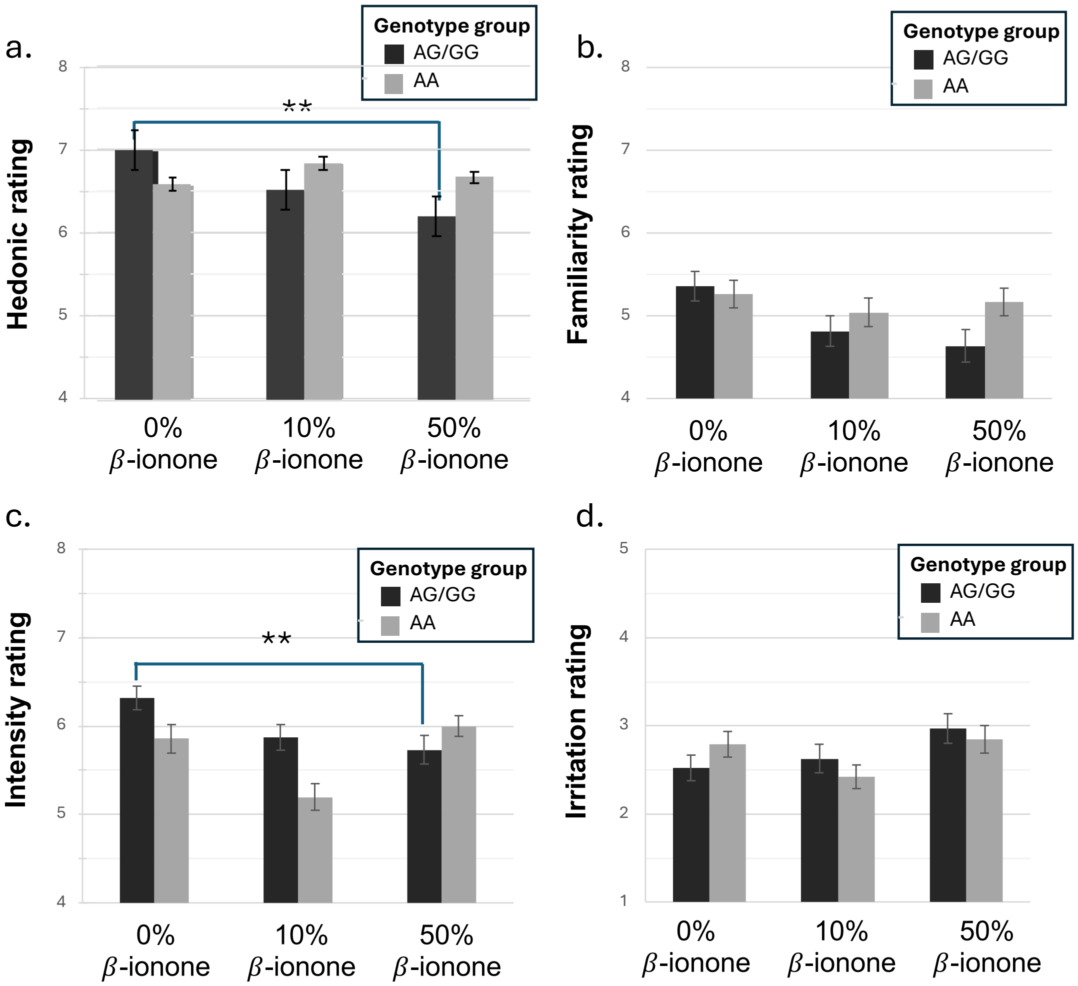


*Figure S3. Number of participants preferring each of the three perfumes depending on genotype group separated by experimental location. a. All three locations combined (n = 168). b. Toulouse location (n = 69). c. Lyon location (n = 29). d. Reims location (n = 70).*

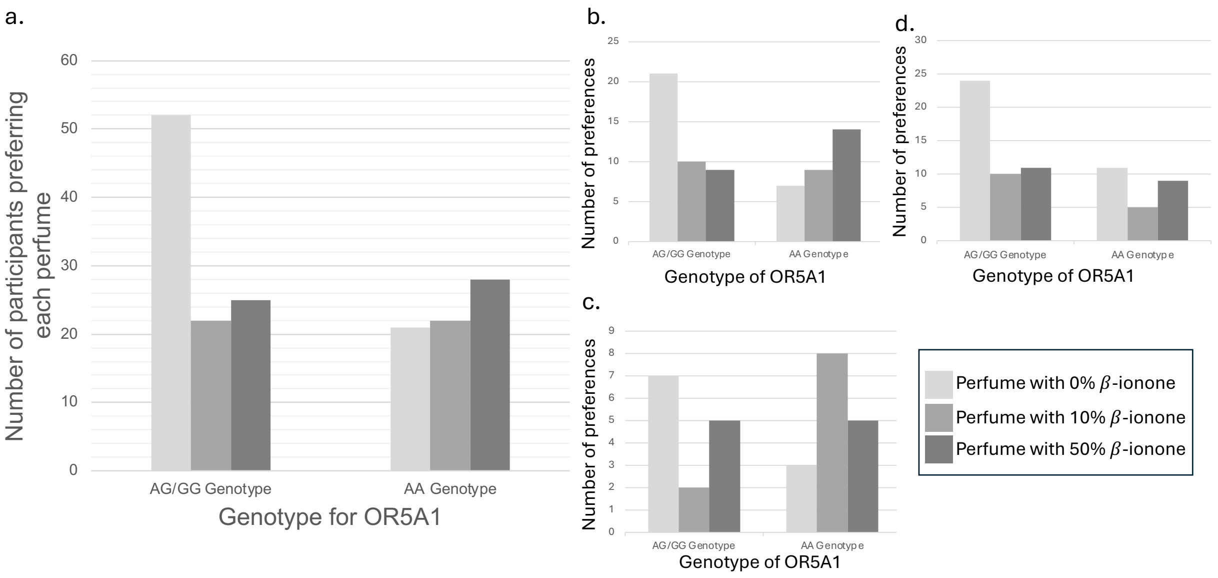
